# Fast and accurate long-read assembly with wtdbg2

**DOI:** 10.1101/530972

**Authors:** Jue Ruan, Heng Li

## Abstract

Existing long-read assemblers require tens of thousands of CPU hours to assemble a human genome and are being outpaced by sequencing technologies in terms of both throughput and cost. We developed a novel long-read assembler wtdbg2 that, for human data, is tens of times faster than published tools while achieving comparable contiguity and accuracy. It represents a significant algorithmic advance and paves the way for population-scale long-read assembly in future.

---

*De novo* sequence assembly reconstructs a sample genome from relatively short sequence reads. It is essential to the study of new species and structural genomic changes that often fail mapping-based analysis as the reference genome may lack the regions of interest. With the rapid advances in single-molecule sequencing technologies by Pacific Biosciences (PacBio) and Oxford Nanopore Technologies (ONT), we are able to sequence reads of 10–100 kilobases (kb) at affordable cost. Such long reads resolve major repeat classes in primates and help to improve the contiguity of assemblies. Nowadays, long-read assembly has become a routine for bacteria and small genomes, thanks to the development of several high-quality assemblers^1-5^. For mammalian genomes, however, existing assemblers may take several CPU years, or ∼1000 US dollars if we use the Google or the Amazon cloud for computing. This price is more expensive than data produced with one PromethION machine, which is capable of sequencing a human genome at 30-fold coverage in two days^6^. To address this issue, we developed wtdbg2, a new long-read assembler that is tens of times faster for large genomes with little compromise on the assembly quality.

Wtdbg2 broadly follows the overlap-layout-consensus paradigm. It advances the existing assemblers with a fast all-vs-all read alignment implementation and a novel layout algorithm based on fuzzy-Bruijn graph (FBG).

For mammalian genomes, current read overlappers^7-9^ split input reads into many smaller batches and perform all-vs-all alignment between batches. This strategy wastes compute time on repeated file I/O and on indexing and querying non-informative *k*-mers. These overlappers do not build a single hash table in worry that it may take too much memory. Interestingly, this should not be a major concern. Wtdbg2 first loads all reads into memory and counts *k*-mer occurrences. It then bins every 256bp on reads into one unit and builds a hash table with keys being *k*-mers occurring twice or more, and values being identifiers of associated bins. For the CHM1 human genome sequenced to 60-fold coverage^10^, for example, there are only 1.5 billion non-unqiue *k*-mers with the default settings. Staging raw read sequences in memory and constructing the hash table takes 250GB at the peak, which is comparable to the memory usage of short-read assemblers.

Wtdbg2 bins read sequences to speed up the next step in alignment: dynamic programming (DP). With 256bp binning, the DP matrix is 65536 (=256×256) times smaller than a per-base DP matrix. This reduces DP to a much smaller scale in comparison to *k*-mer based^8^,^9^ or base-level DP^7^.

FBG extends the basic ideas behind de Bruijn graph (DBG) to work with long noisy reads. In analogy to DBG, a “base” in FBG is a 256bp bin and a “*K*-mer” or *K*-bin in FBG consists of *K* consecutive bins on reads. A vertex in FBG is a *K*-bin and an edge between two vertices indicates their adjacency on a read. Unlike DBG, different *K*-bins may be represented by a single vertex if they are aligned together based on all-vs-all read alignment. This treatment tolerates errors in noisy long reads. FBG is closer to sparse DBG^11^ than standard DBG in that it does not inspect every *K*-bin on reads. The sparsity reduces the memory to construct FBG. Furthermore, FBG explicitly keeps identifiers of bins going through each edge to retain long-range information without a separate “read threading” step as with standard DBG assembly. After graph simplification, wtdbg2 writes the final FBG to disk with read sequences on edges contained in the file. Wtdbg2 constructs the final consensus with partial order alignment^12^ over edge sequences.

We evaluated wtdbg2 v2.3 on five datasets along with CANU-1.8^4^, FALCON-180831^1^, Flye-2.3.6^5^ and MECAT-180314^3^ (Table 1). We used minimap2 to align assembled contigs to the reference genome and to collect metrics. On all datasets, wtdbg2 is at least 4 times as fast as the closest competitors. Wtdbg2 assemblies sometimes cover less reference genomes, which is a weakness of wtdbg2, but its contigs tend to have fewer duplicates. The fraction of genomic regions covered by one contig is only a couple of percent lower in comparison to other assemblers.

**Table 1.**
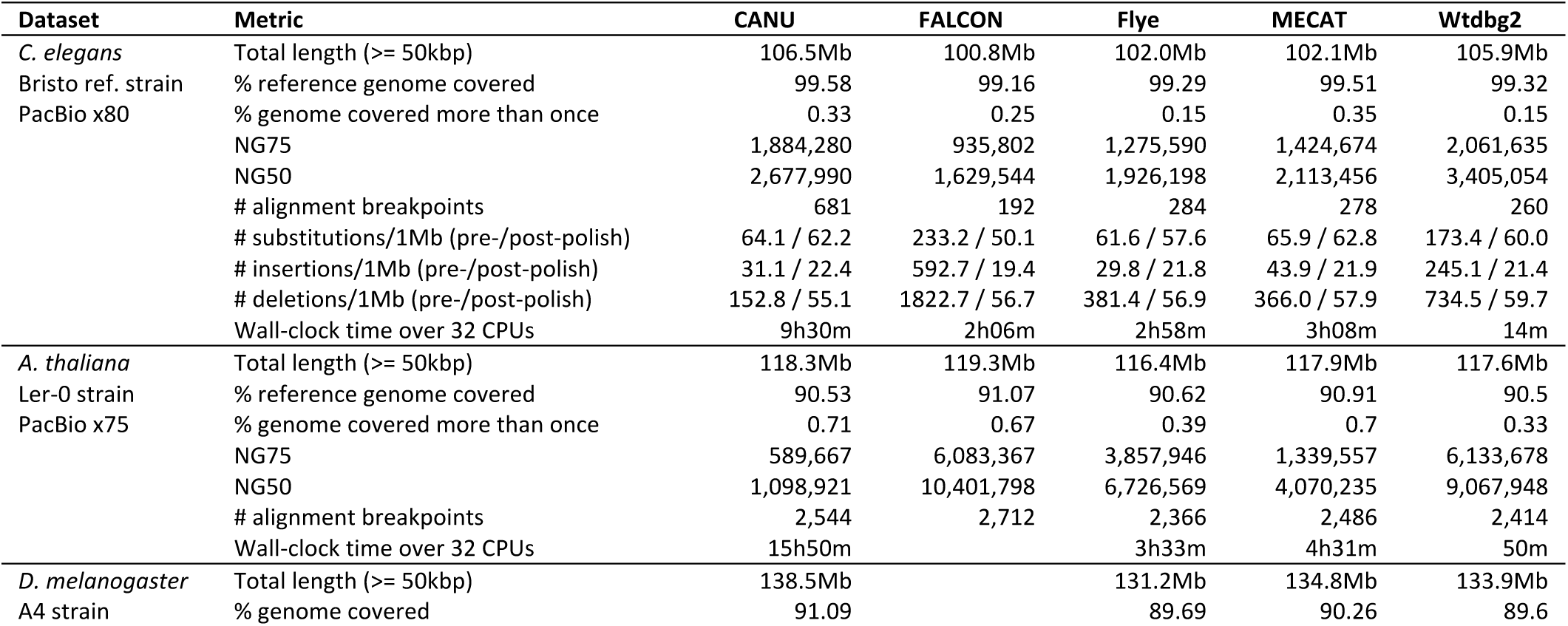
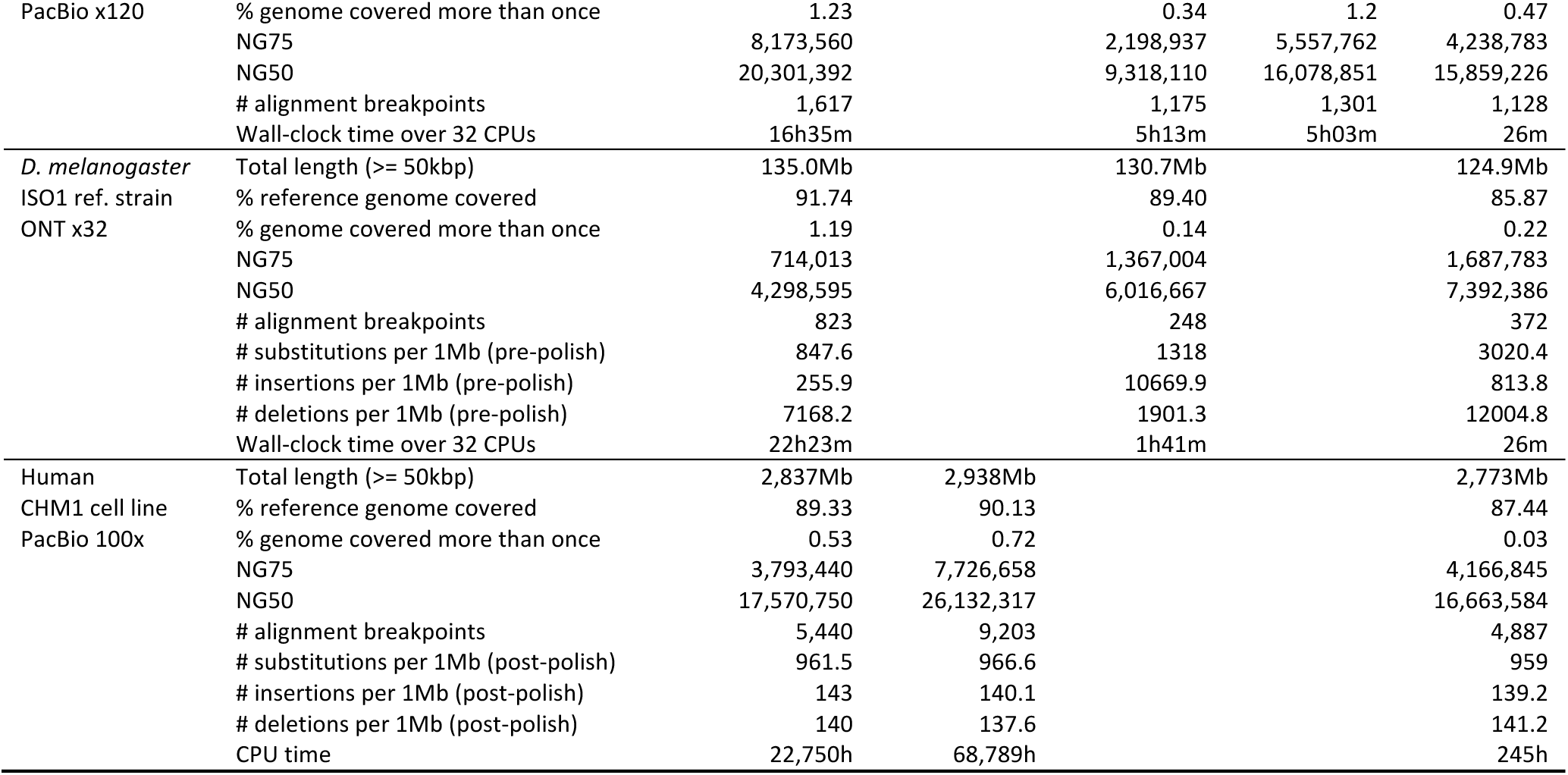
Evaluating long-read assemblies FALCON requires PacBio-style read names and does not work with ONT data or the A4 strain of *D. melanogaster* which was downloaded from SRA. The *A. thaliana* assembly by FALCON is acquired from PacBio website as our assembly is fragmented. MECAT produces fragmented assemblies for the ONT dataset. Human assemblies were performed by the developers of each assembler.

For samples close to the reference genome, we also compared the consensus accuracy before and after signal-based polishing^13^ when applicable. Without polishing, CANU, Flye and MECAT tend to produce better consensus sequences. This is probably because they perform at least two rounds of error correction or the consensus step, while wtdbg2 applies one round of consensus only. After Quiver polishing, the consensus accuracy of all assemblers is very close and significantly higher than the accuracy of consensus without polishing. This observation reconfirms that polishing consensus is still necessary^14^ and suggests that the pre-polishing consensus accuracy is not obviously correlated with post-polishing accuracy.

We assembled five additional large datasets with wtdbg2 (Table 2). For all human data, wtdbg2 finishes the assembly in a few days on a single computer. This performance broadly matches the throughput of a PromethION machine. In comparison, Flye and CANU reportedly required ∼10,000 and ∼40,000 CPU hours, respectively, to assemble NA12878^15^. The peak memory used by wtdbg2 is generally below 230GB, except for HG00733 which was sequenced to a much higher coverage. Wtdbg2 is able to assemble axolotl, the largest genome we have sequenced so far, at a speed 30 times faster than the published assembly^16^ and with a longer NG50.

**Table 2.** Wtdbg2 performance on large genomes

| Data set | Technology | G. size | Cov. | CPU hour | Real hour | NG50 |
| --- | --- | --- | --- | --- | --- | --- |
| Human NA12878 | Nanopore | 3Gb | 36 | 891 | 31 | 11.7Mb |
| Human NA19240 | Nanopore | 3Gb | 35 | 1025 | 35 | 4.3Mb |
| Human NA24385 | PacBio CCS | 3Gb | 28 | 290 | 10 | 12.9Mb |
| Human HG00733 | PacBio Sequel | 3Gb | 93 | 2267 | 52 | 25.9Mb |
| Axolotl | PacBio RSII | 32Gb | 32 | 4262 | 110 | 392kb |

Ten years ago when the Illumina sequencing technology entered the market, the sheer volume of data effectively decommissioned all aligners and assemblers developed earlier. History repeats itself. Affordable population-scale long-read sequencing is on the horizon. Wtdbg2 is the only assembler that is able to keep up with the throughput and the cost. With heterozygote-aware consensus algorithms and phased assembly planned for future, wtdbg2 and upcoming tools might fundamentally change the current practices on sequence data analysis.

**Fig 1.**
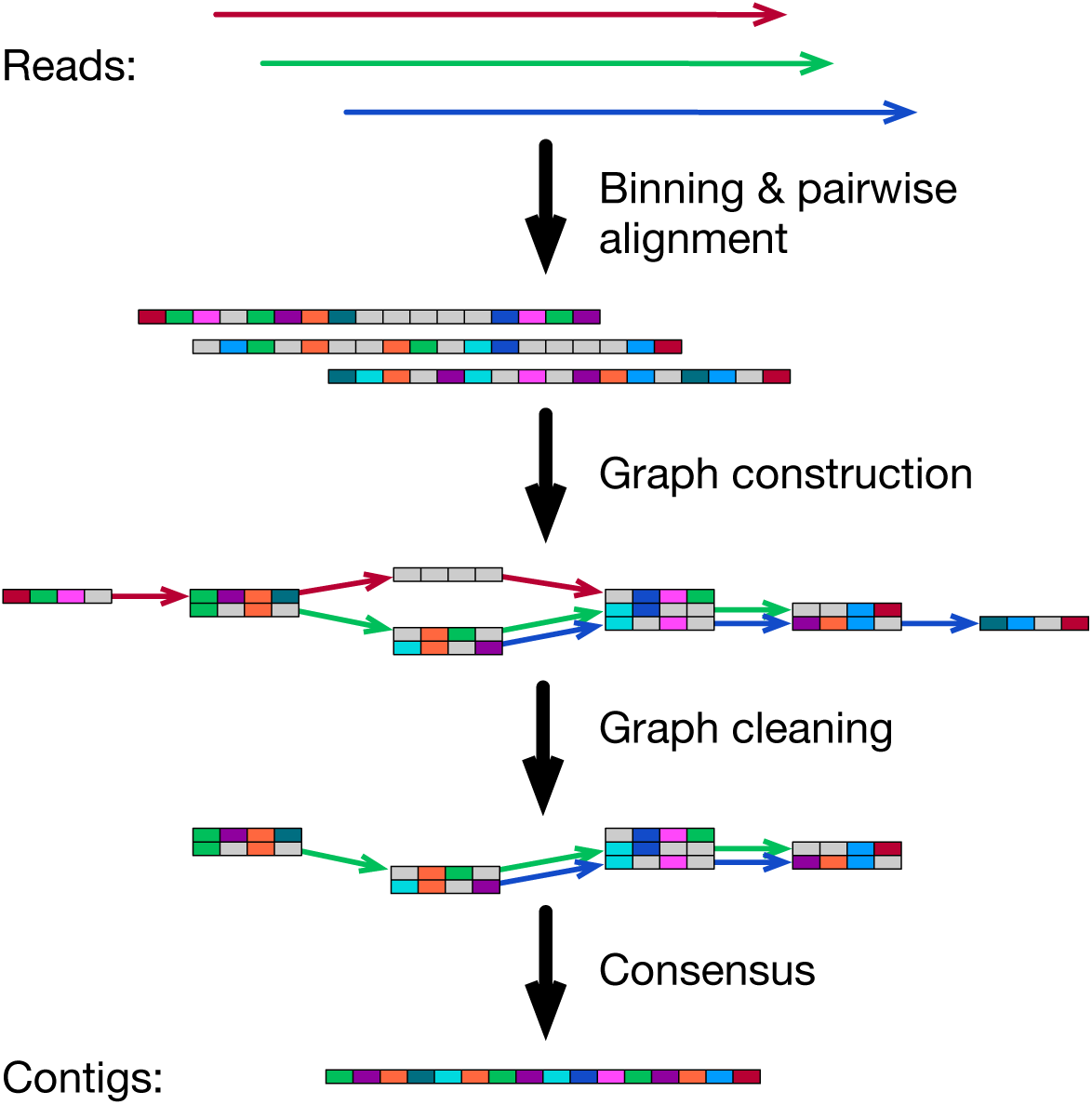
The wtdbg2 algorithm. Wtdbg2 groups 256 base pairs into a bin, a small box in the figure. Bins/boxes with the same color suggest they share *k*-mers, except that a gray bin doesn’t match other bins due to sequencing errors. Wtdbg2 performs all-vs-all alignment between binned reads, ignoring detailed base sequences. In the fuzzy-Bruijn assembly graph, a vertex is a 4-bin segment. Wtdbg2 identifies equivalent segments based on their alignments and connects these segments if they are connected on the input reads (see Online Methods for details). Information on reads going through each edge is kept in the graph (colored edges). After graph construction, wtdbg2 trims tips and pops bubbles and produces the final contig sequences from the consensus of read subsequences attached to each edge.

## Acknowledgements

We would thank Chengxi Ye from University of Maryland for frequent and fruitful discussion in the development of wtdbg. This study was supported by Natural Science Foundation of China (NSFC; grant 31571353 and 31822029 to J.R.) and by US National Institutes Health (NIH; grant R01-HG010040 to H.L.).

## Author contribution

J.R. conceived the project, designed the algorithm and implemented wtdbg2. H.L. contributed to the development and drafted the manuscript. Both authors evaluated the results and revised the manuscript.

## Online methods

### Evaluation datasets

*C. elegans* and *A. thaliana* reads are provided by PacBio at http://bit.ly/pbpubdat. We downloaded SRR5439404 for the *D. melanogaster* A4 strain, SRR6702603 for the *D. melanogaster* reference ISO1 strain, PRJNA378970 for axolotl, SRR7615963 for HG00733, and ERR2631600 and ERR2631601 for NA19240. CHM1 reads were acquired from http://bit.ly/chm1p6c4, NA12878 reads from http://bit.ly/na12878ont (release 5) and NA24385 from http://bit.ly/NA24385ccs.

### Wtdbg2 algorithm

Wtdbg2 reads all input sequences into memory and encodes each base with 2 bits. By default, it selects a quarter of *k*-mers based on their hash code and counts their occurrences using a hash table with 46-bit key to store a *k*-mer and 17-bit value to store its count. Wtdbg2 filters out *k*-mer occurring once or over 1000 times in reads, and then scans reads again to build a hash table for the remaining *k*-mers and their positions in bins.

For all-vs-all read alignment, wtdbg2 traverses each read, from the longest to the shortest, and uses the hash table to retrieve the reads that share *k*-mers with the read in query. It applies Smith-Waterman-like DP between binned sequences, penalizing gaps and bins that do not share *k*-mers. Wtdbg2 retains alignments no shorter than 8×256bp. After finishing alignments for all reads, wtdbg2 frees the hash table but keeps the all-vs-all alignments in memory (alignments are also written to disk as intermediate results).

At this step, wtdbg2 loses base sequences. It only sees binned sequences and the alignments between them. On a binned sequence *B* = *b*_1_*b*_2_ … *b*_|*B*|_, a *K*-bin *b*_*Ki*_ = *b*_*i*_*b*_*i*+1_ … *b*_*i*+*K*-1_ is a *K*-long subsequence starting at *i*-th position on *B*. If binned sequences *B* and *B*’ can be aligned, we can infer the overlap length between *K*-bins *b*_*Ki*_ and 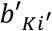 by lifting their coordinates between the two sequences based on the alignment. We say two *K*-bins are equivalent if they completely overlap. Using the all-vs-all alignment, wtdbg2 collects a non-redundant set Ω of *K*-bins such that no *K*-bin in Ω is equivalent to others. Wtdbg2 also records the locations and coverage of each *K*-bin in Ω.

The vertex set *V* of FBG is a subset of Ω. To construct *V*, wtdbg2 traverses each non-redundant *K*-bin in the descending order of their original coverage. Given a *K*-bin *b*_*K*_, wtdbg2 may reduce its coverage by deducting *K*-bins in *V* that overlap with *b*_*K*_. If the adjusted coverage is ≥3 and higher than half of the original coverage, *b*_*K*_ will be added to *V*; otherwise it will be ignored. After the construction of *V*, wtdbg2 adds an edge between two *K*-bins if they are located on the same read. There are often multiple edges between two *K*-bins. Wtdbg2 retains one edge and keeps the count. An edge covered by <3 reads are discarded. This generates FBG. The coverage thresholds can be adjusted on the wtdbg2 command line.

### Assembling evaluation datasets

With wtdbg2, we specified the genome size and sequence technology on the command line, which automatically applies multiple options. Specifically, we used “-xrs -g100m” for *C. elegans*, “-xsq -g125m” for *A. thaliana*, “-xrs - g144m” for *D. melanogaster* A4 strain, “-xont -g144m” for the ISO1 strain, “-xrs -g3g” for CHM1, “-xont -g3g” for human NA12878 and NA19240 ONT reads, “-xsq -g3g” for HG00733, “-xccs -g3g” for NA24385 and “-xrs -g3g” for the axolotl dataset. We used the similar settings for CANU, Flye and MECAT, specifying only the genome size and the sequencing technology. The FALCON configure file for assembling *C. elegans* is provided as **supplementary data**. The FALCON *A. thaliana* assembly was downloaded at http://bit.ly/pbpubdat. We are using AC:GCA_000983455.1 for the CANU CHM1 assembly and AC:GCA_001297185.1 for the FALCON CHM1 assembly.

### Evaluating assemblies

To count alignment breakpoints, we mapped all assemblies to the corresponding reference genomes with minimap2 under the option “--paf-no-hit - cxasm20 -r2k -z1000,500”. We used the companion script paftools.js to collect various metrics (command line: “paftools.js asmstat -q50000 -d.1”). To count substitutions and gaps, we applied a different minimap2 setting “-cxasm5 --cs -r2k”. This setting introduces more alignment breakpoints but avoids poorly aligned regions harboring spuriously high number of differences that are likely caused by large-scale variations and skew the counts. We used “paftools.js call” to call variations.

### Code availability

The wtdbg2 source code is hosted by GitHub at: https://github.com/ruanjue/wtdbg2.

### Data availability

All the evaluated assemblies (except those publicly available in GenBank) can be obtained at ftp://ftp.dfci.harvard.edu/pub/hli/wtdbg/. The FTP site also provides the detailed command lines and configuration files.

